# Neuronal EphB2 signaling drives persistent neuropathic pain following spinal cord injury

**DOI:** 10.64898/2026.04.20.719620

**Authors:** Nicolette M. Heinsinger, David A. Jaffe, Kolluru D. Srikanth, Megan A. Lyttle, Madison S. Smith, Samantha J. Thomas, Brittany A. Charsar, Lan Cheng, Pauline Michel-Flutot, Rachel E. Cain, Jaime L. Watson, Duran Bao, Jia Fan, Aditi Falnikar, Wei Zhou, Matthew B. Dalva, Angelo C. Lepore

## Abstract

Neuropathic pain after spinal cord injury reflects persistent hyperexcitability in the spinal cord dorsal horn, yet the molecular drivers sustaining this maladaptive state are unknown. Using an antibody microarray of dorsal horn tissue from mice six weeks after cervical contusion spinal cord injury, we found persistent upregulation of Eph-ephrin signaling, including increased EphB1, EphB2 and EphB3 expression and phosphorylation. Reversible chemogenetic inhibition of EphB kinase activity, using an EphB1/2/3 analog-sensitive knock-in mouse, selectively reversed established mechanical allodynia without affecting thermal hyperalgesia or motor function and also shifted dorsal horn signaling away from pain sensitization-associated pathways. Among EphB receptors, EphB2 showed the most consistent and robust injury-induced increase in expression within dorsal horn. Although EphB2 transcript levels increased in both dorsal horn neurons and astrocytes, conditional deletion of EphB2 only in dorsal horn neurons, but not in astrocytes, reversed established mechanical allodynia and reduced dorsal horn neuronal activation. These findings identify EphB signaling, and neuronal EphB2 in particular, as a mechanism that actively maintains pain hypersensitivity after spinal cord injury.

## INTRODUCTION

Neuropathic pain is among the most debilitating consequences of spinal cord injury and is often refractory to pharmacological treatment.^1^ Chronic pain after spinal cord injury can manifest as hyperalgesia, allodynia, or spontaneous pain, each reflecting maladaptive sensory processing within the dorsal horn.^2–4^ Despite extensive study and therapeutic efforts, only a minority of patients achieve meaningful relief with current medications, and reversal of established neuropathic pain remains rare.^5–8^ Although dorsal horn hyperexcitability is recognized as a central driver of this condition,^2,9^ the molecular mechanisms that maintain this neuropathic pain state, and that might be targeted to reverse pathological pain, remain unknown. This gap is important because mechanisms required for maintenance of pathological pain may differ from those that initiate it, and identifying these mechanisms could reveal targets capable of reversing chronic pain once it is established.

Persistent changes in protein abundance and phosphorylation are well-positioned to sustain long-term shifts in excitability and synaptic plasticity. Among signaling systems with the potential to do so, EphB receptor tyrosine kinases are notable.^10–14^ EphBs regulate excitatory neurotransmission through phosphorylation-dependent interactions with NMDA receptors, activate MAPK and PKC cascades implicated in central sensitization, and have been linked to neuropathic pain after peripheral nerve injury.^10–12,15–18^ However, the functional role played by EphB receptors in spinal cord injury-induced neuropathic pain has not been examined. To identify candidate pathways that remain altered after spinal cord injury, we used an antibody microarray to quantify changes in protein expression and phosphorylation in parallel and found persistent upregulation of EphB signaling.

Here, we tested the hypothesis that persistent neuropathic pain after spinal cord injury is sustained by ongoing EphB-dependent signaling in dorsal horn neurons. We combined unbiased molecular profiling with reversible chemogenetic inhibition of EphB kinase activity in vivo and conditional deletion of EphB2 in defined dorsal horn cell populations to determine whether EphB signaling is required for maintaining chronic pain. These approaches identify EphB signaling, and EphB2 in particular, as a mechanism that actively maintains neuropathic pain after spinal cord injury and suggest that targeting this pathway may restore more normal sensory processing, even after chronic pain has become established.

## RESULTS

### Eph-ephrin signaling was upregulated following cervical spinal cord injury

Unilateral cervical (C5/6) contusion spinal cord injury produces a chronic neuropathic pain-related phenotype characterized by persistent thermal hyperalgesia and mechanical allodynia in the ipsilesional forepaw, as well as spontaneous pain-related behavior.^19–21^ To identify signaling pathways that might sustain this condition, we used a proteomics-based antibody microarray to quantify total and phosphorylated proteins in the ipsilesional dorsal horn (at spinal cord levels C7-C8) six weeks after injury, when behavioral hypersensitivity was fully established.^19,22^ Tissue samples from spinal cord injury and laminectomy-only uninjured control mice were analyzed (Figure 1A-B). The array included 2,059 antibodies (each spotted in duplicate) recognizing 1,165 distinct proteins and 894 phosphorylation sites, enriched for receptor and intracellular signaling pathways (Kinexus Bioinformatics Corporation, Vancouver Canada) (Figure 1C-D).^22^

**Figure 1.**
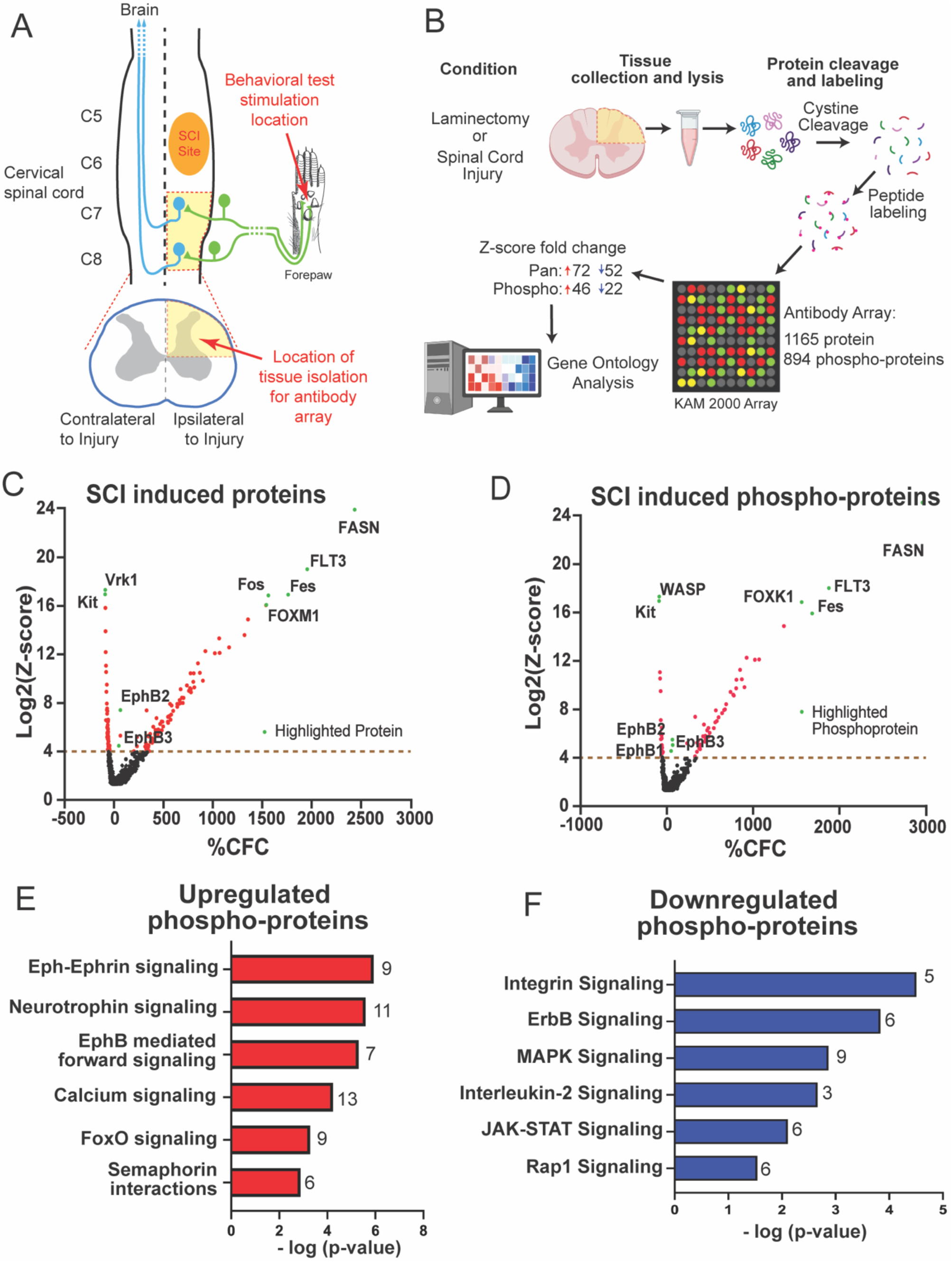
Effects on protein expression and phosphorylation in the dorsal horn after cervical spinal cord injury. (A) Diagram shows location of the unilateral contusion spinal cord injury site at level C5-C6, as well as the C7-C8 location where ipsilateral dorsal quadrant was microdissected to harvest tissue homogenate for antibody array, immunoblotting and qPCR analyses. (B) Diagram of experimental approach. For both experimental groups, three samples were generated by pooling together the dorsal horn quadrants of two animals per sample (6 total mice per group: 2 females pooled; 2 males pooled; 2 males pooled). Arrays assayed 1,165 different proteins and 894 different phosphoration sites from 562 distinct proteins. Protein cleavage, binding, and analysis were conducted by Kinexus (Vancover, Canada). (C-D) Volcano plots depicting the antibody array changes in total proteins and phospho-sites after cervical spinal cord injury. The %CFC (percent change from control sham: laminectomy-only) indicates the mean expression level for each protein or phospho-site. Each dot represents one protein or phospho-site. The black dots represent non-significant differentially expressed proteins or phospho-sites. The red dots indicate significant differentially expressed proteins or phospho-sites, and the green dots indicate a labelled subset. The differentially regulated proteins were determined by using an unpaired t-test. (E-F) Gene Ontology analysis of changes in signaling pathways determined by changes in phospho-proteins following spinal cord injury (contusion) compared to laminectomy-only sham control; (E) shows upregulated pathways, and (F) shows downregulated. Numbers on the end of the bars indicate number of phospho-proteins that significantly changed in response to spinal cord injury.

Spinal cord injury induced extensive remodeling of the dorsal horn proteome, with 74 proteins significantly upregulated and 52 downregulated (log₂ [z-score] > 4; Figure 1C). Phosphoprotein analysis revealed that 46 sites increased and 22 decreased after spinal cord injury relative to uninjured control (Figure 1D). Proteins known to increase neuronal excitability, such as calcium/calmodulin-dependent protein kinase 2, CREB1 and Fos, increased in expression after spinal cord injury (Supplemental File 1).^23–25^ In addition, injury led to increased expression and phosphorylation of expected inflammation-associated proteins such as fatty acid synthase (FASN) and c-Jun (Supplemental File 1).^26–28^ Conversely, proteins known to be anti-inflammatory in certain contexts such as insulin-like growth factor 1 showed decreased expression and phosphorylation in dorsal horn after spinal cord injury (Supplemental File 1).^29^ Among the most consistently upregulated phosphoproteins were EphB1, EphB2 and EphB3, implicating EphB receptor signaling as persistently elevated after injury (Figure 1D).

Gene ontology (GO) enrichment analysis of upregulated proteins in spinal cord injury compared to laminectomy-only mice demonstrated Eph-Ephrin signaling was most strongly upregulated, along with neurotrophin and calcium signaling pathways (Figure 1E). Downregulated pathways included integrin and ERB signaling (Figure 1F). These findings reveal that spinal cord injury drives long-term activation of certain EphB-dependent signaling networks in the dorsal horn – EphB-dependent pathways known to regulate excitatory neurotransmission and plasticity ^17,30,31^ – and suggest that this signaling may contribute to neuropathic pain maintenance.

### Inhibition of EphB tyrosine kinase activity reversed already-established neuropathic pain-related behavior after spinal cord injury

Based on the antibody array data showing that injury-induced neuropathic pain was associated with increased EphB1-3 expression and elevated EphB signaling (Figure 1), we asked whether inhibiting EphB and its downstream signaling pathways might reduce, or even reverse, the neuropathic pain-like phenotype. To test whether EphB kinase signaling is required for maintaining established neuropathic pain in spinal cord injury mice, we used a chemogenetic model, where EphB kinase activity was rendered sensitive to a designer drug, 1-naphthyl-PP1 (1-NA-PP1), by point mutation to the ATP-binding pocket of the intracellular tyrosine kinase domain of EphB1, EphB2, and EphB3 (Figure 2A).^32^ The analog sensitive-EphB triple knock-in (AS-EphB TKI) mouse enables selective, inducible, and reversible blockade of the kinase activity of EphB1, 2, and 3.^32^ Consistent with a model of EphB dysfunction selectively in chronic pain,^18,33,34^ 1-NA-PP1 delivery to uninjured laminectomy-only mice resulted in no behavioral effects on mechanical or thermal sensitivity (Figure 2E, G).

**Figure 2.**
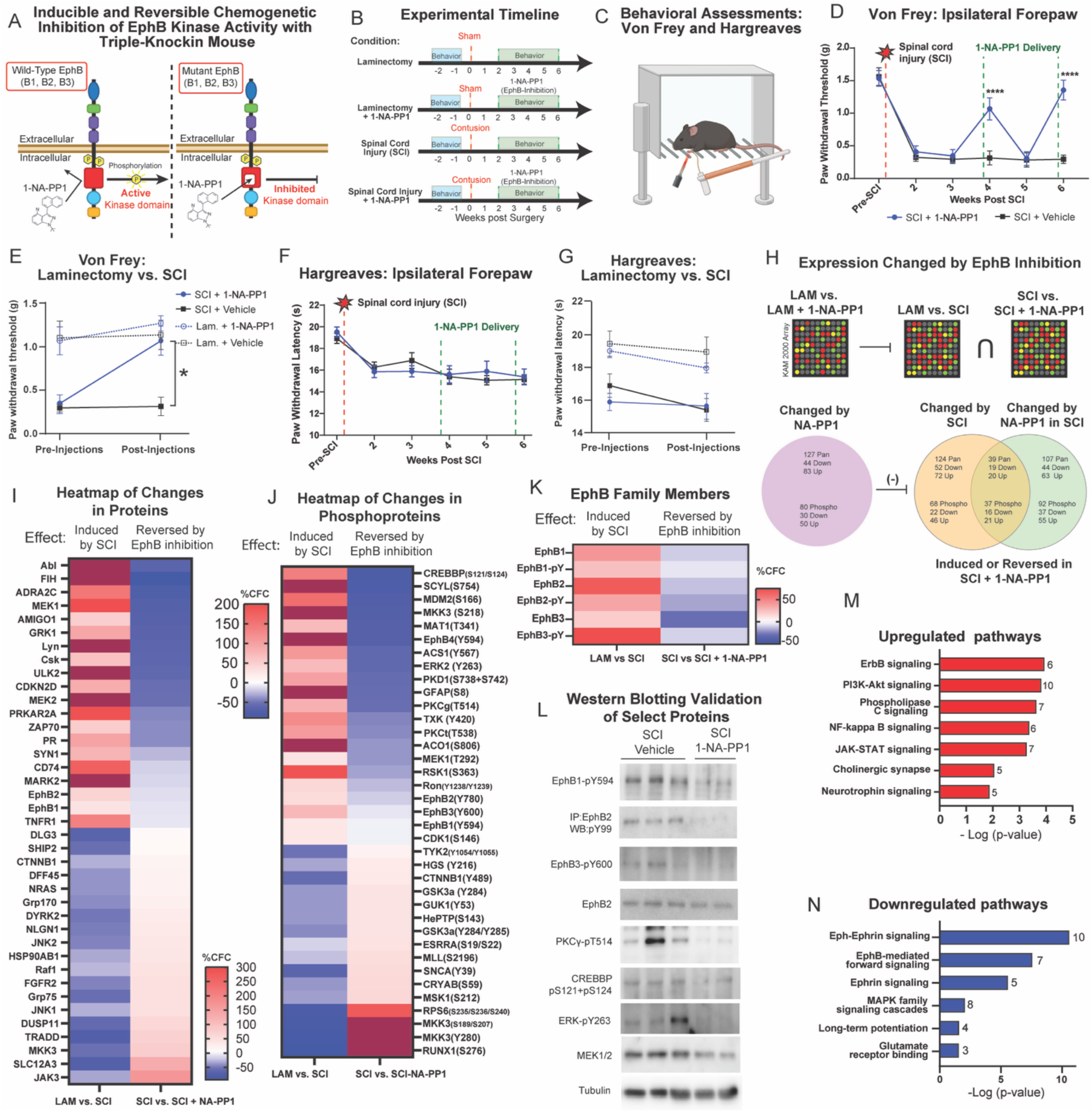
Inducible chemogenetic EphB kinase inhibition reversed mechanical allodynia after spinal cord injury. (A) Model illustrating the experimental approach and the method of EphB kinase inhibition. (B) Timeline of experimental paradigm. (C) Behavioral paradigm for either Von Frey or Hargreaves testing; positive responses were recorded only when accompanied by some form of supraspinal response (licking, guarding, vocalization, etc.). (D-G) Behavioral effects of inhibition of EphB kinase activity. Von Frey test for mechanical sensitivity (D) and Hargreaves test for thermal sensitivity (F) in spinal cord injury mice that were administered either 1-NA-PP1 or a vehicle-only control. Behavioral results for timepoints pre-treatment and post-treatment (with 1-NA-PP1 or vehicle-only) in spinal cord injury and laminectomy-only mice: Von Frey test (E) and Hargreaves test (G). Two-way repeated measures ANOVA was used to assess significance in graphs D-G. D and F, Bonferroni’s and Šídák’s multiple comparisons, respectively, were made between groups at each time point. E and G, Bonferroni’s multiple comparisons were made between groups and over time. * = p<0.05, ** = p<0.01, *** = p<0.001, **** = p<0.0001. (H) Dorsal quadrant of C7-C8 spinal cord from mice was microdissected, and antibody microarray analysis was performed on this tissue. Laminectomy-only AS-EphB TKI mice and spinal cord injury AS-EphB TKI mice that received 1-NA-PP1 treatment were compared to their respective controls that received vehicle-only treatment. For both of the spinal cord injury groups, four samples were generated by pooling together the dorsal horn quadrants of two animals per sample (8 total mice per group: 2 females pooled; 2 females pooled; 2 males pooled; 2 males pooled). For both of the laminectomy-only groups, three samples were generated by pooling together the dorsal horn quadrants of two animals per sample (6 total mice per group: 2 females pooled; 2 males pooled; 2 males pooled). (I) Heatmap of proteins that show differential regulation in spinal cord injury versus laminectomy-only (column 1) compared to spinal cord injury treated with 1-NA-PP1 versus vehicle control (column 2). Proteins changed significantly following injury and blockade of EphB kinase activity with 1-NA-PP1. (J) As in I, a heat map of phosphorylation sites on the indicated proteins is shown. (K) Effects of inducible EphB kinase inhibition on expression and phosphorylation of EphB1, EphB2, and EphB3. (L) Western blot validation of select proteins and phosphoproteins that are part of neuronal pathways involved in chronic pain. (M-N) Gene Ontology analysis of up- and down-regulated pathways in response to spinal cord injury following inducible EphB kinase inhibition. Numbers on the end of the bars indicate number of phospho-proteins that significantly changed. Detailed statistical analysis for this figure is provided in Supplemental Table 2.

To test the impact of inducible inhibition of EphB1-3 kinase activity on neuropathic pain-related behavior following spinal cord injury, baseline sensory behavioral measurements were assessed, cervical contusion spinal cord injury was then delivered, and follow-up weekly behavioral testing was performed (Figure 2B-C). Four weeks after spinal cord injury when neuropathic pain is well-established,^19,20,35^ three doses of vehicle-only control or 1-NA-PP1 were injected intraperitoneally every 12 hours. Neither vehicle control nor 1-NA-PP1 had any effect on thermal hyperalgesia (Hargreaves, Figure 2F-G; Supplemental Figure 1B). However, injection of 1-NA-PP1, but not vehicle control, robustly reversed established forepaw mechanical allodynia bilaterally (Von Frey filament, Figure 2D-E; Supplemental Figure 1A). Consistent with the importance of EphB signaling for the maintenance of long-lasting mechanical hypersensitivity, the effects of EphB inhibition were lost one day after stopping 1-NA-PP1 administration (Figure 2D; Supplemental Figure 1A). A subsequent series of 1-NA-PP1 injections, two weeks after the initial dosing series, produced a similar reversal of mechanical allodynia (Figure 2D; Supplemental Figure 1A). The effects of inhibiting EphB kinase activity were selective to sensory behavior, as no differences were observed between vehicle-only and 1-NA-PP1-treated AS-EphB TKI mice in grip strength in either forepaw (Supplemental Figure 1C-D). These results demonstrate that EphB kinase activity is required to maintain persistent mechanical allodynia after CNS damage.

### Inducible EphB tyrosine kinase inhibition reduced activation of downstream pathways associated with dorsal horn neuron excitability following spinal cord injury

Along with reversal of established mechanical allodynia in spinal cord injury mice (Figure 2D-E; Supplemental Figure 1A), inducible inhibition of EphB kinase activity is likely to result in downstream changes in protein expression and phosphorylation in dorsal horn. To determine the molecular profile associated with this behavioral outcome, protein array changes in the ipsilesional dorsal horn (Figure 1A-B) were determined in mice following spinal cord injury and blockade of EphB kinase activity using 1-NA-PP1. This analysis was performed in four groups of animals: 1) laminectomy-only AS-EphB TKI mice with vehicle-only control injection; 2) laminectomy-only AS-EphB TKI mice with 1-NA-PP1 injection; 3) spinal cord injury AS-EphB TKI mice with vehicle-only; and 4) spinal cord injury AS-EphB TKI mice with 1-NA-PP1. The behavioral effects of 1-NA-PP1 were validated in AS-EphB TKI mice by assessing the reversal of spinal cord injury-induced mechanical hypersensitivity.

Laminectomy-only AS-EphB TKI mice were given 1-NA-PP1 to control for effects of the small molecule independent of SCI. Proteins and phosphorylation sites that changed significantly after 1-NA-PP1 treatment in uninjured mice were removed from further analysis (Figure 2H). After controlling for effects of 1-NA-PP1, analysis of antibody microarrays revealed that 1-NA-PP1 reversal of allodynia was associated with increased expression of 79 proteins and decreased expression of 40 proteins. Additionally, 45 phosphorylation sites were increased, and 27 were decreased. As expected, EphB1-3 were among those proteins with significantly decreased phosphorylation (Figure 2K; Supplemental File 2). These data suggest that inhibition of EphB kinase activity after spinal cord injury drives dorsal horn proteome changes that may be associated with reduced hypersensitivity.

Next, we sought to determine whether blocking EphB signaling results in changes in protein expression and phosphorylation that are altered by the neuropathic pain state. A simple model for the role of EphB signaling in maintaining neuropathic pain would suggest that proteins and phosphorylation sites that increase or decrease after spinal cord injury would be changed in the opposite direction following EphB kinase inhibition and ensuing reversal of mechanical hypersensitivity. A total of 39 proteins and 37 phosphorylation sites followed this pattern (Figure 2I-J; Supplemental File 2). Consistent with this model, proteins downregulated after spinal cord injury in response to EphB inhibition include PKC, CREBBP, Mek1, Mek2, and Erk (Figure 2I); inhibition of these proteins is known to reduce chronic pain.^36–39^ The antibody array data for a number of proteins impacted by EphB inhibition after spinal cord injury were validated by immunoblotting, including the effects on EphB1-3 phosphorylation (Figure 2L). Notably, some proteins previously associated with chronic pain like JNK1, JNK2, and MKK3 were decreased after spinal cord injury, but increased with EphB inhibition (Figure 2I).^40,41^ Collectively, these data suggest that EphB-dependent signaling regulates multiple sets of proteins in the dorsal horn whose expression and signaling are involved in neuropathic pain.

To begin to define the molecular signature of EphB kinase-dependent neuropathic pain pathways, antibody array data on phospho-protein changes were subjected to GO analysis. GO enrichment and pathway analysis identified several signaling pathways that were suppressed by spinal cord injury and re-engaged upon EphB inhibition. Notably, proteins associated with ErbB receptor, PI3K-Akt, phospholipase-C, and NF-κB signaling showed significant enrichment among those upregulated in EphB-inhibited spinal cord injury samples (Figure 2M). These results indicate that blocking EphB kinase activity weeks after injury, when long-lasting allodynia has already been established, can restore or enhance multiple pro-growth and cell-survival pathways initially dampened by injury.^42–45^ Conversely, pathways known to drive maladaptive pain were less active when EphB signaling was blocked; GO analysis indicated reduced enrichment of long-term potentiation and MAPK cascade pathways in EphB-inhibited spinal cord injury mice relative to the uninjured condition (Figure 2N). These pathways are known to contribute to central sensitization and chronic pain states.^46,47^ Taken together, these data demonstrate that inducible EphB kinase inhibition after spinal cord injury causes a broad shift in downstream signaling networks, down-regulating pain-facilitating pathways while up-regulating pathways involved in repair and normal synaptic function. Such molecular alterations likely underlie the robust behavioral improvement in neuropathic pain-like behavior after spinal cord injury observed with EphB inhibition, and these data place EphB receptors as a regulatory hub, influencing both adaptive and maladaptive aspects of spinal cord plasticity.

### EphB2 gene and protein expression increased following cervical spinal cord injury

We further assessed whether spinal cord injury alters expression levels of the three EphBs targeted with the AS-EphB TKI mice, EphB1-3. When quantitative PCR was performed on the dorsal cervical spinal cord of uninjured animals, *Ephb1* mRNA was found to be expressed at approximately four times the amount of *Ephb2*, while *Ephb3* was expressed at about half the level of *Ephb2* (Figure 3A). We next tested two cervical spinal cord injury paradigms, unilateral C5/6 contusion with 40 kilodyne (moderate) or 60 kilodyne (severe) impact force. To examine whether there were long-lasting changes in EphB expression, animals were allowed to survive for six weeks post-injury, and then EphB expression was determined using qPCR of the ipsilateral dorsal horn quadrant (Figure 3E). Animals were assessed for mechanical allodynia and thermal hyperalgesia prior to sacrifice to validate that the mice were in a neuropathic pain-like state. Quantification of qPCR indicated that, regardless of impact force, *Ephb1* mRNA expression did not change in the dorsal spinal cord 6 weeks post-injury (Figure 3B). *Ephb3* mRNA levels increased after 60 kilodyne spinal cord injury, but not with the less severe 40 kilodyne contusion (Figure 3D). In contrast, *Ephb2* expression significantly increased after both 40 and 60 kilodyne spinal cord injury (Figure 3C). The increased expression of *Ephb2* and *Ephb3*, combined with the lack of a change in *Ephb1* mRNA levels after injury, suggest that these two members of the EphB family might be particularly associated with the persistent pain state.

**Figure 3.**
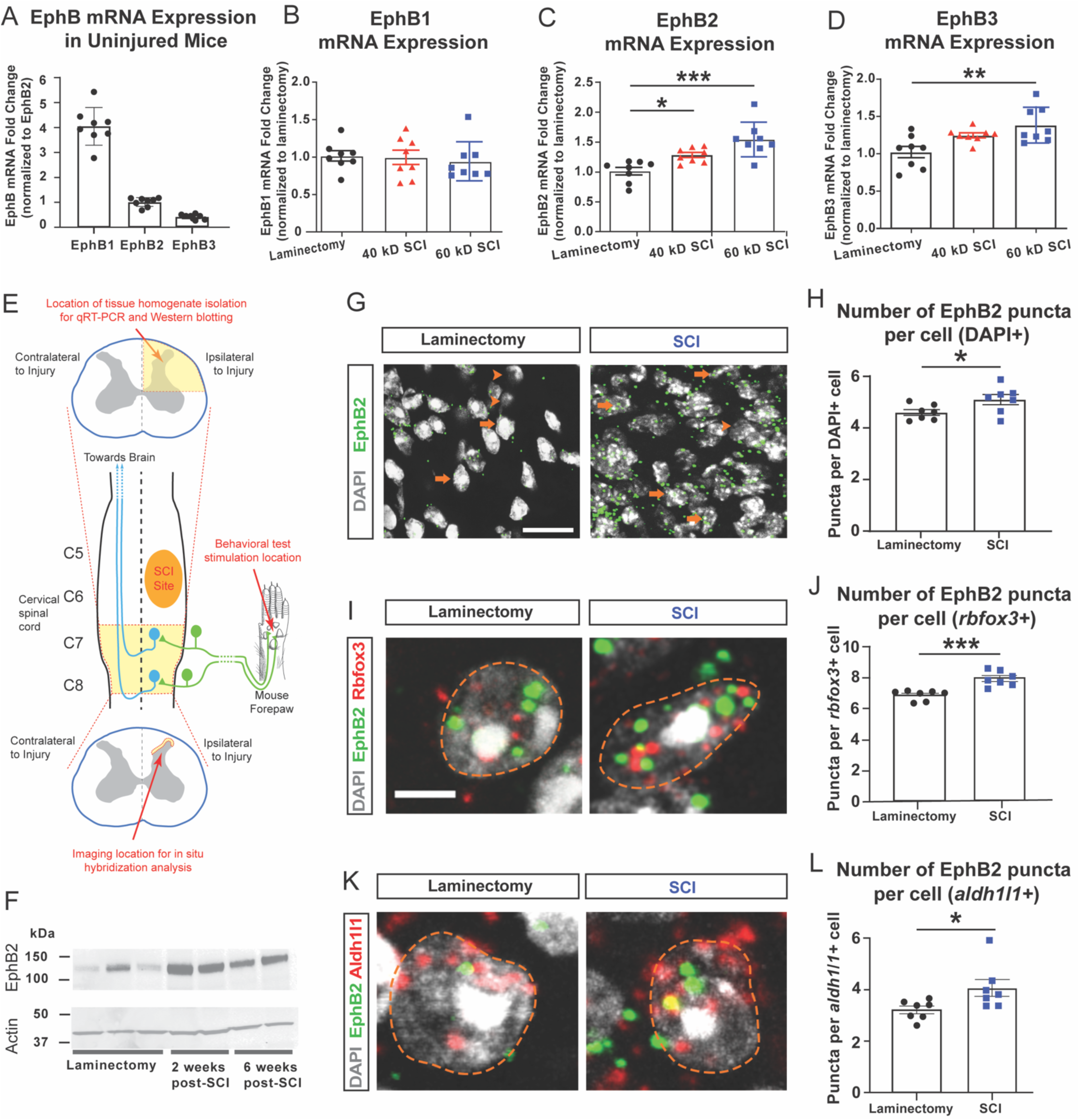
EphB2 expression was increased in neurons and astrocytes of cervical superficial dorsal horn after spinal cord injury. (A) qPCR quantification of *Ephb1*, *Ephb2,* and *Ephb3* mRNA expression in the cervical dorsal horn of uninjured spinal cord, normalized to *EphB2* level. qPCR quantification of (B) *Ephb1*, (C) *Ephb2*, and (D) *Ephb3* in the cervical dorsal horn of laminectomy-only versus spinal cord injury mice. (E) Diagram showing location of the contusion site at C5/C6, as well as the superficial dorsal horn location in caudal ipsilesional spinal cord where RNAscope and immunohistochemistry analyses were performed. (F) Immunoblotting comparing spinal cord tissue from animals undergoing laminectomy-only procedure versus mice at 2 and 6 weeks after spinal cord injury. RNAscope *in situ* hybridization was used to quantify *Ephb2* expression in neurons and astrocytes after spinal cord injury. Representative images of *Ephb2* mRNA expression in the superficial dorsal horn two weeks after either (G, I, K – left) laminectomy-only or (G, I, K – right) spinal cord injury. (G) Arrowheads indicate cells with no *Ephb2* expression, and arrows indicate cells with *Ephb2* expression. (I) Representative images of *Ephb2* expression in *Rbfox3*+ neurons. Orange line outlines the DAPI+ nucleus. (J) Quantification of *Ephb2* puncta per *Rbfox3*+ neuron. (K) Representative images of *Ephb2* expression in *Aldh1l1*+ astrocytes. (L) Quantification of *Ephb2* puncta per *Aldh1l1*+ astrocyte. One-way ANOVA using Tukey’s multiple comparisons test was performed to analyze differences in graphs A-D. * = p<0.05, ** = p<0.01, *** = p<0.001, **** = p<0.0001. An independent (unpaired) samples t-test was used to determine significance with * = p<0.05, ** = p<0.01, *** = p<0.001 compared to laminectomy-only value for graphs H, J, and L. Detailed statistical analysis for this figure is found in Supplemental Table 3.

To determine in which dorsal horn cell types EphB2 or EphB3 expression changes occur, RNAscope *in situ* hybridization was conducted for *Ephb2* and *Ephb3* in laminectomy-only and cervical contusion mice in ipsilesional C7 spinal cord just caudal to the injury site. This analysis was performed specifically in superficial dorsal horn laminae, given the importance to pain transmission of this location in dorsal horn (Figure 3E).^48–51^ There were no differences in *Ephb3* expression between laminectomy-only uninjured and spinal cord injury conditions in superficial dorsal horn across all DAPI+ cells or specifically in *Rbfox3*+ neurons (Supplemental Figure 2A-D). Consistent with the qPCR and protein data from dorsal spinal cord whole-tissue homogenate, *Ephb2* mRNA expression in superficial dorsal horn increased compared to uninjured control (Figure 3G-H). The number of *Ephb2* transcripts increased after spinal cord injury both in *Rbfox3*+ neurons (Figure 3I-J) and in *Aldh1l1+* astrocyte (Figure 3K-L).

Of the three EphB family proteins, EphB2 expression in dorsal horn appeared to be most sensitive to injury. Therefore, to validate whether changes in mRNA levels extended to protein expression changes, immunoblotting of the dorsal spinal cord of laminectomy-only and 40 kilodyne cervical contusion was performed. These data demonstrate a persistent upregulation of EphB2 protein levels ipsilateral to the cervical spinal cord injury both at 2 weeks and 6-weeks post-contusion (Figure 3F). Together with the qPCR, RNAscope and protein array data, these findings demonstrate that EphB2 is upregulated in dorsal horn by spinal cord injury and suggest that EphB2 may be linked to long-lasting pain hypersensitivity.

### EphB2 knockout in neurons of cervical dorsal horn reversed already-established mechanical allodynia

To determine whether the persistent increase in dorsal horn EphB2 expression contributes to maintenance of neuropathic pain-like behavior, *Ephb2* was conditionally deleted from dorsal horn neurons or dorsal horn astrocytes after mechanical allodynia was established. Floxed-EphB2 mice underwent contusion and, after allowing for hypersensitivity to become chronic with a five-week recovery period, mice received focal intraspinal microinjections of AAV-hSynapsin-Cre or AAV-GFAP-Cre into the ipsilesional superficial dorsal horn (at segments C7-C8) to induce *Ephb2* knockout selectively in neurons or astrocytes, respectively (Figure 4A-C). This design allowed repeated assessment of pain-like behavior after EphB2 deletion at 2, 4, and 6 weeks post-injection (7, 9, and 11 weeks after injury), allowing us to test whether neuronal EphB2 is required for the maintenance of established mechanical hypersensitivity.

**Figure 4.**
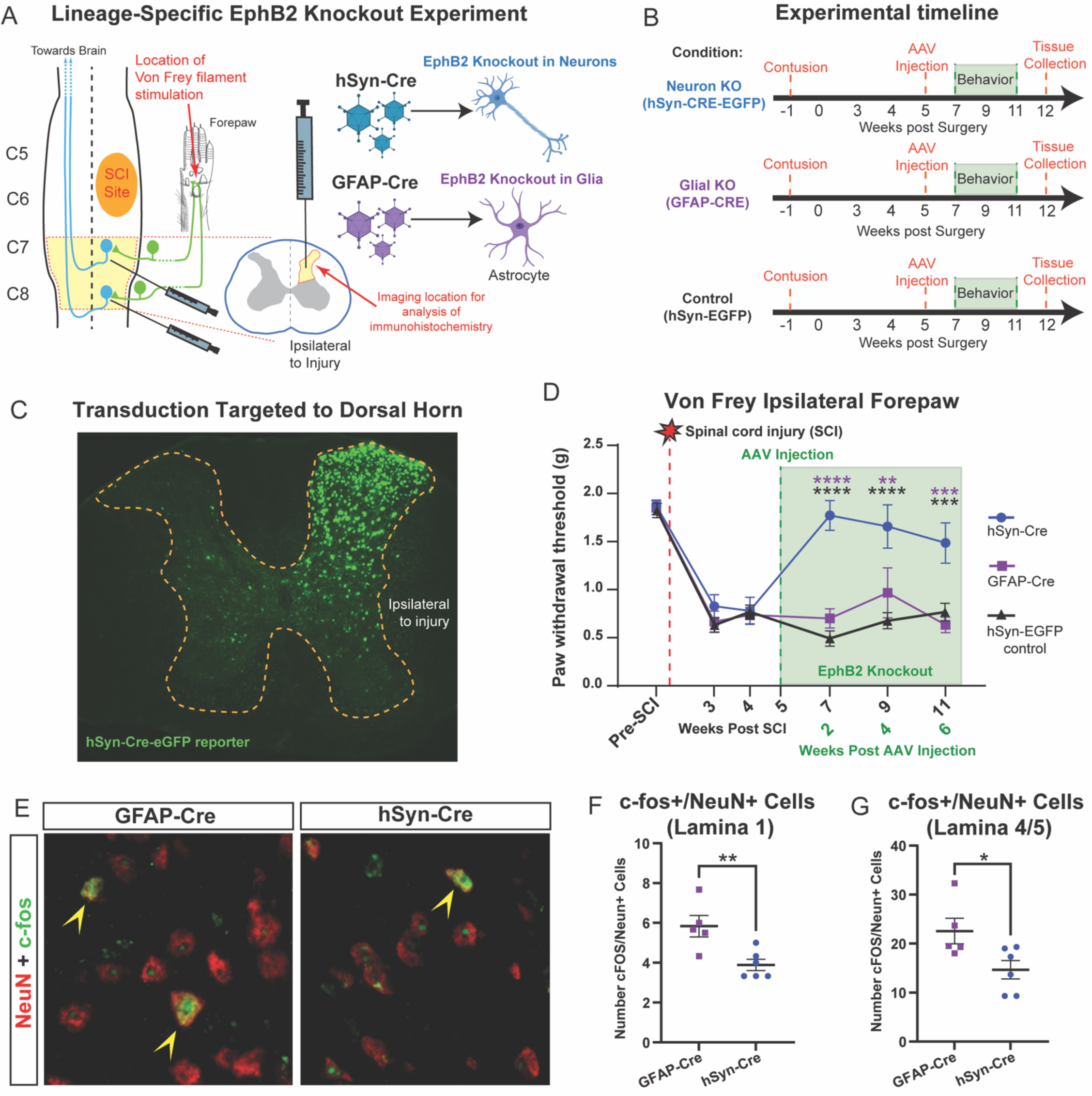
Inducible neuron-specific *Ephb2* knockout in cervical dorsal horn following spinal cord injury reversed already established mechanical allodynia. (A) Diagram depicting the location of the C5/C6 contusion injury, as well as the ipsilesional C7 and C8 location for AAV microinjection and immunohistochemistry analysis. The diagram also includes the model for neuron versus astrocyte targeted lineage-specific *Ephb2* knockout. (B) Timeline of behavioral experiments relative to contusion injury and viral injections. (C) Representative image of eGFP reporter of AAV transduction targeted to the ipsilesional dorsal horn. (D) Von Frey test quantification, comparing mice with Cre driven by either a human synapsin (hSyn-Cre) promoter or glial fibrillary acidic protein (GFAP-Cre) promoter, along with a third AAV group where no Cre is expressed (hSyn-EGFP). (E) Immunohistochemistry was used to label c-Fos positive cells and NeuN positive neurons; example images for (E, left) GFAP-Cre and (E, right) hSyn-Cre are shown. Cells showing co-labeling of c-Fos and NeuN are marked with an arrowhead. (F-G) Quantification of the number of cells co-labeled with both c-Fos and NeuN in (F) lamina 1 and (G) lamina 4/5 in mice injected with either GFAP-Cre or hSyn-Cre vectors. In panel D, two-way ANOVA using Bonferroni’s multiple comparisons were performed made between groups and over time. In panels F and G, an independent (unpaired) samples t-test was used to determine significance between groups. For all graphs, * = p<0.05, ** = p<0.01, *** = p<0.001 indicate significance between groups. Detailed statistical analysis for this figure is provided in Supplemental Table 4.

AAV-mediated recombination was both efficient and cell-type specific in the dorsal horn: 87.9 +/- 1.4% of NeuN-positive neurons were transduced by AAV-hSynapsin-Cre-GFP, and 88.9 +/- 1.2% of GFP reporter-positive transduced cells were NeuN-positive. Control animals receiving AAV-hSynapsin-GFP (no Cre vector) maintained stable hypersensitivity across all time points (Figure 4D). In addition, mechanical sensitivity was not affected by targeting of *Ephb2* knockout to glia with AAV-GFAP-Cre injection (Figure 4D). In contrast, neuron-specific knockout of *Ephb2* in dorsal horn led to a significant and long-lasting reversal of mechanical allodynia after AAV-hSynapsin-Cre-GFP injection (out to at least 11 weeks post-injury, Figure 4D). Consistent with our findings that EphB2 acts selectively on hypersensitivity after spinal cord injury, inducible *Ephb2* knockout had no effect on forelimb motor function as assessed by grip strength (Supplemental Fig. 3A). These findings indicate that EphB2 expression in dorsal horn neurons, but not neighboring glia, is required to maintain established mechanical hypersensitivity after spinal cord injury.

Neuron-specific *Ephb2* knockout reversed already-established mechanical sensitivity in spinal cord injury mice. To determine whether this robust behavioral effect was associated with changes in neuronal activation, we measured c-Fos levels in dorsal horn neurons. While the total number of NeuN-positive cells in dorsal horn was similar across groups (Supplemental Figure 3C-D), neuron-specific *Ephb2* knockout significantly reduced numbers of c-Fos expressing neurons in ipsilesional C7-C8 dorsal horn in both superficial lamina 1 (Figure 4E-F) and in deeper laminae 4-5 (Figure 4G), but not in laminae 2-3 (Supplemental Figure 3B). Together, these data indicate that EphB2 expression in dorsal horn neurons is required for the maintenance of mechanical hypersensitivity and is also associated with increased dorsal horn neuronal activation after spinal cord injury.

## DISCUSSION

How persistent changes in signaling pathways sustain maladaptive behavior is a central question in neuroscience. Neuropathic pain after spinal cord injury provides a clear example: dorsal horn hyperexcitability persists long after the initial insult, yet the molecular mechanisms that actively maintain this state remain poorly defined.^1,52–54^ Our findings identify EphB signaling, and EphB2 in particular, as one important mechanism. Three observations support this conclusion. First, unbiased protein array analysis revealed persistent upregulation of Eph-ephrin signaling weeks after injury. Second, reversible inhibition of EphB kinase activity reduced established mechanical allodynia after hypersensitivity was already present. Third, conditional deletion of EphB2 selectively in dorsal horn neurons, but not in astrocytes, reversed established mechanical hypersensitivity. Together, these findings indicate that EphB signaling is not simply induced by injury, but remains continuously required to sustain the sensitized state.

The protein array data further suggest that EphB signaling sits upstream of multiple pathways implicated in chronic pain. Spinal cord injury produced persistent changes in both protein abundance and phosphorylation, extending prior work beyond transcriptional changes alone and identifying a molecular signature more closely linked to ongoing signaling state. Inhibition of EphB kinase activity after injury suppressed MAPK- and long-term potentiation-associated pathways, while re-engaging signaling networks linked to growth and survival. This bidirectional shift suggests that EphB signaling helps stabilize a broader maladaptive molecular configuration in the dorsal horn, rather than acting through a single downstream effector. EphB receptors are well positioned to do so, given their known ability to regulate kinase cascades such as MAPK and PKC ^10,16,34^ and to influence synaptic plasticity through phosphorylation-dependent mechanisms.^12–14^

At the same time, the effects of EphB inhibition were selective. Blocking EphB kinase activity reversed mechanical allodynia but did not affect thermal hyperalgesia, and inducible EphB2 deletion also had no effect on forelimb motor function. In addition, the requirement for EphB2 localized to dorsal horn neurons near the injury site, despite the broader distribution of pain-processing circuits. These findings argue that EphB-dependent signaling does not function as a general regulator of sensory processing, but instead supports defined mechanisms underlying established mechanical hypersensitivity after injury. Whether EphB2 plays similar roles in other neuronal populations, anatomical locations, pain modalities, and injury models remains to be determined.

More broadly, the properties of EphB2 distinguish it from many candidate pain mechanisms. EphB2 expression and signaling is low in the uninjured spinal cord, significantly increases after neural injury, and remains required after chronic hypersensitivity is established. EphB2 therefore emerges not simply as part of the acute injury response, but as a signal required to maintain the chronic pain state. The ability to impact established hypersensitivity by inhibiting EphB signaling suggests that the chronic state remains reversible because it continues to depend on ongoing signaling rather than circuit changes alone.

## Resource availability

All unique/stable reagents generated in this study are available from the lead contacts with a completed materials transfer agreement. The lead contact for the mice is MBD.

## Acknowledgements

This work was supported by NINDS (R01NS110385 to ACL, MBD; R01NS079702 to ACL) and the Yant Family Spinal Cord Regeneration Fund (to ACL). We thank Michael Greenberg and Henry Ho for EphB-TKI mice.

## Author contributions

Conceptualization: ACL, MBD

Methodology: NMH, DAJ, KDS, MAL, ACL, MBD

Investigation: NMH, DAJ, KDS, MSS, SJT, BAC, LC, PMF, REC, JLW, DB, JF, AF, WZ

Visualization: NMH, DAJ, KDS

Funding acquisition: ACL, MBD

Project administration: ACL, MBD

Supervision: ACL, MBD

Writing – original draft: NMH, DAJ, KDS, ACL, MBD

Writing – review & editing: DAJ, KDS, ACL, MBD

## Declaration of generative AI and AI-assisted technologies

Generative AI tools were used to check grammar and assist in sentence flow.

## Supplemental information titles and legends

Figures S1-S3, Tables S1-S7, Files S1-2

## STAR METHODS

### Experimental model and study participant details

#### Mice

All animal procedures and standard of care were approved by the Thomas Jefferson University Institutional Animal Care and Use Committee (IACUC) and were also carried out according to ARRIVE *(Animal Research: Reporting of* In Vivo *Experiments)* guidelines. Procedures and animal care also followed the NIH Guide for the Care and Use of Laboratory Animals. Three types of mice were used in the experiments: wild-type C57BL/6J mice; AS-EphB2 TKI (EphB1^T697G^, EphB2^T699A^, EphB3^T706A^) with C57BL/6J background; and floxed-EphB2 mice (EphB2^flox/flox^) with C57BL/6J background. All mice were kept in ventilated isolator cages in temperature and humidity-controlled rooms (23°C and 40%-60% humidity). Animals were maintained on a 12-hour light/dark cycle (6:00am start time) and given free access to autoclaved water and food pellets (5010 LabDiet, MN). All experiments utilized both C57BL/6 male (28-35g) and female (22-30g) mice aged 3-5 months old.

### Method details

#### Cervical contusion spinal cord injury

Mice were given a unilateral C5/6 contusion spinal cord injury, similar to previously described work.^55^ Animals were first anesthetized with 4% isoflurane supplied into an anesthesia box, then given 0.05mg/kg buprenorphine via subcutaneous injection. Once animals were unresponsive to bilateral toe pinches, isoflurane was supplied via a nose cone and maintained at 1.5-2.0% depending on animal weight and breathing rate. An incision was made from cervical level 3 (C3) to thoracic level 1 (T1), and then the skin and muscles overlaying the spinal column were retracted. Laminectomies of the vertebrae overlying C5 and C6 were conducted to gain access to the spinal cord. Spinal cord injury mice received a right-sided contusion injury using the Infinite Horizon Impactor (Precision Systems and Instrumentation; Lexington, KY) with an impactor tip of 0.7 mm in diameter, a force of either 40 or 60 kilodynes, and a 2 second dwell time. Uninjured control animals received the same laminectomy with no contusion injury to the spinal cord.

#### Viral vectors

Three different vectors were used: pAAV.GFAP.Cre.WPRE.hGH (Addgene, MA; Catalog # 105550-AAV5), pAAV.hSyn.EGFP (Addgene, MA; Catalog #50465-AAV5), and AAV.hSyn.HI.eGFP-Cre.WPRE.SV40 (Addgene, MA; Catalog # 105553-AAV5), with an AAV5 serotype. Cre recombinase was driven by either a glial fibrillary acidic protein (GFAP) promoter to direct expression to astrocytes or a human synapsin (hSyn) promoter to drive expression in neurons. A control virus, expressing no Cre, was used with an hSyn promotor to target EGFP expression in neurons. Once received, all virus were thawed on ice, then distributed into 5μL aliquots before freezing at -80C until intraspinal injection use.

#### Intraspinal injection

Ephb2^flox/flox^ mice received unilateral injections of AAV (freshly thawed on ice) at C7 and C8 using similar methods as other studies.^56–58^ In a anesthesia box, animals received 4% isoflurane until fully anesthetized, and then were switched to 1.5-2.0% isoflurane via nose cone. Skin and all muscle layers over C3-T1 were retracted to expose the spinal column. A laminectomy above C7 and C8 was conducted to expose the spinal cord, and then the dura was carefully incised at C7 and C8. Only the right side of the animal was injected (ipsilesional to the contusion injury) using a 34G needle attached to a 10-µL Hamilton syringe. Injections were 0.5µL in volume each, 0.2 mm deep, 1 mm lateral to the spinal midline, and delivered at a speed of 0.1µL/min.

#### Behavioral testing

Grip strength, Hargreaves thermal, and von Frey filament testing were performed as previously described on both male and female mice.^21^ These procedures are described below. Before behavioral testing, all mice were acclimated to the researcher and to the room for 1-2 hours per day for two weeks before baseline testing and again for 30-60 mins before each day of testing.

#### Grip strength testing

The DFIS-2 Series Digital Force Gauge (Ametek, PA) was used to measure grip strength.^57^ All measurements were recorded in grams. Mice were lightly scruffed, and one forepaw was placed on the metal pull bar. Experimenters ensured that mice formed a proper grasp around the bar before proceeding. With their head positioned down at a 45-degree angle, mice were gently pulled away horizontally until the paw released the pull bar. Six baseline measurements were collected per paw, and, on every day of testing after baseline, three measurements were collected. There was at least one minute in between each trial.

#### Hargreaves thermal testing

Hargreaves testing was performed on a glass surface with an infrared heat source placed underneath (UgoBasile, VA).^59^ Mice were lightly scruffed with their forepaw placed on the glass just above the heat source. Experimenters ensured the plantar surface of the forepaw was directly over the heat source. The latency to withdrawal of the forepaw in response to the thermal stimulation was measured, with a maximum cut-off time of 30s. Only movement away from the heat source with some form of behavior indicative of a supraspinal response (vocalization, paw flapping, licking, etc.) was recorded. Otherwise, the trial was repeated after a two-minute waiting period. Three trials were conducted per paw per testing day, with at least two minutes in between each trial.

#### Von Frey mechanical testing

Animals were placed in acrylic chambers, without bottoms, atop a metal grate. Semmes-Weinstein monofilaments (Stoetling Company, IL) were used to assess mechanical sensitivity using the von Frey up-down method.^21,59^ Filaments were applied to the center of the plantar surface of the forepaw until the filament bent; if a positive withdrawal response was observed, the next smaller filament was used, and if no withdrawal response occurred then the next higher filament was used. Ten trials were performed per paw, starting with the 0.6g filament, with 2 minutes in between each trial. A maximum cut-off was set at the 2g filament. Forepaw withdrawal was considered a quick removal of the paw from the stimulus, which was also accompanied by an indication of supraspinal awareness of the stimulus such as licking or flapping of the forepaw, guarding of the forepaw, vocalization, etc. Movements in which the researchers did not observe a supraspinal response were not recorded, and the trial was repeated.

#### Spinal cord dissection

Animals were sacrificed after spinal cord injury or laminectomy-only by anesthetic overdose of ketamine-xylazine cocktail. Animals used for immunohistochemistry were then perfused with 0.9% saline, followed by 4% paraformaldehyde. The spinal cord was then harvested, fixed in 4% paraformaldehyde at 4°C overnight, and washed in 0.1M phosphate buffer at 4°C for 24 hours. Samples were then cryoprotected in a 30% sucrose solution at 4°C for three days. Tissue used for RNAscope was prepared fresh frozen without use of paraformaldehyde. For both fresh frozen and fixed frozen tissue, C3-T1 region was removed from the rest of the spinal cord, embedded in tissue freezing medium (General Data, Cincinnati, Ohio), and flash frozen in 2- methylbutane (Fisher Scientific, Pittsburgh, Pennsylvania). Tissue was cut transversely by a cryostat into 30 µm thick sections and placed directly onto glass slides (Fisher Scientific, Pittsburgh, Pennsylvania). Sectioned tissue was stored at -20⁰ C until use.

#### Western blotting

Immediately after animal sacrifice, ipsilateral dorsal quadrant several segments caudal to injury was sub-dissected and flash frozen in liquid nitrogen. Samples were separately stored at -80°C until use. Tissue samples were thawed and then homogenized on ice in 50 µL of RIPA buffer (50 mM TRIS-HCl pH 7.6, 150 mM NaCl, 2 mM EDTA, 0.1% SDS, 0.01% NP-40, and Protease Inhibitor Cocktail; Roche Diagnostics, Indianapolis, IN). Equal amounts of protein, as determined by Bradford assay, were run on 10-12% Bis-Tris SDS-PAGE gels and transferred to nitrocellulose membranes. Membranes were blocked at room temperature for one hour with 5% milk (when testing proteins) or 2% BSA (when testing phospho-proteins) mixed in Odyssey blocking buffer (Li-Cor, Lincoln, NE). Membranes were probed with primary antibodies (Table 1) at 4°C overnight. This was followed by HRP-conjugated anti-goat/anti-rabbit/anti-mouse secondary (1:10,000; Jackson Immunoresearch, West Grove, PA) or IRDye-conjugated goat anti-mouse IgG (1∶20,000; Li-Cor, Lincoln, NE) at room temperature for one hour. Biorad gel doc/Li-Cor Odyssey infrared imaging system was used to image membranes.

#### EphB2 western blotting

The immunoblotting protocol used has been previously described.^9^ Animals were sacrificed two or six weeks after contusion or laminectomy-only via anesthetic overdose and perfused with 0.9% saline. The cervical spinal cord was dissected and flash frozen in dimethylbutane. Ipsilateral dorsal quadrant at the level of injury or several segments caudal to the injury (C7-C8) was sub-dissected, and these samples were separately stored at -80°C until use. Tissue samples were homogenized on ice in 50 µL of RIPA buffer (50 mM TRIS-HCl pH 7.6, 150 mM NaCl, 2 mM EDTA, 0.1% SDS, 0.01% NP-40, and Protease Inhibitor Cocktail) (Roche Diagnostics, Indianapolis, IN). Equal amounts of protein, as determined by the Bradford assay, were run on 4-12% Bis-Tris SDS-PAGE gels and transferred to nitrocellulose membranes. Membranes were blocked at room temperature for one hour with Odyssey blocking buffer (Li-Cor, Lincoln, NE). Membranes were probed for EphB2 (1:500; R&D Systems, Minneapolis, MN) and actin (1∶2,000; Abcam, Cambridge, UK) at 4°C overnight, followed by HRP-conjugated anti-goat secondary (1:10,000; Jackson ImmunoResearch, West Grove, PA) or IRDye-conjugated goat anti-mouse IgG (1∶20,000; Li-Cor, Lincoln, NE) at room temperature for one hour. Li-Cor Odyssey infrared imaging system was used to image membranes, and band intensity was measured and normalized to actin intensity via ImageJ software. All values were then normalized to laminectomy-only levels.

#### Co-immunoprecipitation of phospho-EphB2

Immunoprecipitation followed by western blotting was necessary to assess phospho-EphB2 levels. Spinal cords of AS-EphB2 TKI mice, with either 1-NA-PP1 subcutaneous administration or with vehicle-only control, were homogenized and lysed using RIPA buffer containing protease inhibitors and agitated at 4°C. Tissue lysates were harvested and centrifuged at 13,000 rpm for 25 minutes to pellet down debris. An additional Percoll gradient spin was carried out following lysis to remove myelin and unwanted cellular debris. Proteins were quantified, and an equal amount of protein was used to perform immunoprecipitation. The supernatant was incubated with antibody goat polyclonal anti-EphB2 (Cat# AF467, R&D Systems, Minneapolis, MN) to conjugate on ice for 2 hrs. Antibody-bound proteins were then isolated using pre-blocked protein-G agarose beads (Invitrogen, Waltham, MA) on a rotator at 4 °C. Samples were centrifuged and beads were washed four times in RIPA lysis buffer and two times in TBS-V. Immunoprecipitants were eluted from the agarose beads by adding boiling SDS-sample buffer and boiled at 95 °C. Lysates were separated using 8% SDS-polyacrylamide gels and transferred onto 0.45 µm PVDF membranes (Millipore, Burlington, MA). Immunoblots were then blocked in 5% nonfat dry milk or 2% Bovine Serum Albumin in TBS-T (150 mM NaCl, 10 mM Tris pH 8.0, 0.05% Tween-20). Primary antibodies were presented in blocking solution for 2 hrs at room temperature or overnight at 4°C. HRP-conjugated secondary antibodies were used at 1:10,000 in blocking solution for 1 hr (Jackson ImmunoResearch, West Grove, PA) and then visualized using ECL (PerkinElmer, Waltham, MA) at Biorad Imager.

#### RNA isolation and quantitative real time PCR

The ipsilateral dorsal quadrant of C7 and C8 was microdissected and homogenized in Trizol (Invitrogen, Waltham, MA) to extract mRNA, according to the manufacturer’s instructions. Chloroform was added and tubes were shaken rigorously until phase separation. Tubes were centrifuged at 12,000 x g for 15 mins at 4°C, and then the aqueous phase was removed and placed in a new tube. RNA was precipitated using isopropanol at -20°C overnight and then centrifuged at 12,000 x g for 20 mins. The supernatant was removed, and the pellet was washed with 75% ethanol. The pellet was centrifuged at 7,5000 x g for 5 mins, the ethanol removed, and the pellet dried. cDNA was synthesized using a total of 20 µl of RNA, Superscript IV (Invitrogen, Waltham, MA) and oligo (dT)12–18 (Invitrogen, Waltham, MA), plus 1 µl of RNase H (Invitrogen, Waltham, MA). Tubes were incubated for 20min at 37°C. RT-PCR reactions were run on the 7500 Real Time PCR System (Applied Biosystems, Waltham, MA) using SYBR green PCR master mix (Applied Biosystems, Waltham, MA), 6.35ng cDNA, and 0.5μM forward and reverse primer mix. *Hprt* was used as an internal control, and all samples were normalized to *Hprt* levels. Three sample triplicates were averaged together for each analysis.

#### RNAscope

RNAscope *in situ* hybridization multiplex version 2 was performed on fresh frozen spinal cord sections according to instructions from Advanced Cell Diagnostic (ACD Bio, Newark, CA).^60^ Tissue was fixed using RNAse-free 4% PFA at room temperature for 1 hour and then dehydrated with 50%, 70%, 100% and another 100% ethanol for 5 mins each. Hydrogen peroxide was applied to the slides for 10 mins and then washed with 1x PBS. Protease IV reagent was then applied for 30 mins and washed with 1xPBS. Channel 1, Channel 2, and Channel 3 probes were mixed at a ratio of 50:1:1 and incubated in the HybEZ oven (ACD Bio, Newark, CA) for 2 hours at 40°C. Slides were washed in 1x RNAscope wash buffer twice and then incubated with Amp1 (30 mins), Amp2 (30 mins), and Amp3 (30mins). Slides were washed twice in between each incubation step. Slides were then incubated with HRP-C1 (15 mins), an Opal fluorophore (520, 570, or 690; Akoya Biosciences, Marlborough, MA) (30 mins), and HRP blocker (15 mins), with two washes between each step. The HRP and fluorophore steps were repeated for Channel 2 and Channel 3. The following mouse probes were used: EphB2-C1 (447611), EphB3-C2 (510251-C2), Rbfox3-C2 (313311-C2), Aldh1l1-C3 (405891-C3). Images were taken with a confocal microscope (Leica SP8) using a 40x objective. Three 100µm x 100µm sections were imaged across three spinal cord slices per animal, and the average number of puncta per DAPI-stained nuclei across all images was measured per animal.

#### Immunohistochemsitry

Antibody staining was completed similarly to our previous work.^55,61^ Briefly, slides with cervical spinal cord tissue slices were placed in PBS for three 5-minute washes to remove excess freezing medium. Slides were then left in solution consisting of 5% normal donkey serum (NDS) and 0.2% triton in PBS for 30 minutes at 23°C. After another PBS wash, slides were soaked for 180 minutes at 23°C in primary antibody solution (5% NDS and 0.1% Triton). c-Fos was labelled using a rat anti-c-Fos primary antibody at a 1:500 dilution (Synaptic Systems – Göttingen, Germany; Catalog #226017). NeuN labelling was accomplished with a rabbit anti-NeuN primary antibody at a 1:500 dilution (Cell Signaling Technology, MA; Catalog # D4G4O). After PBS washing, slides were left in secondary antibody solution (5% NDS and 0.1%Triton) for 120 minutes at 23°C. Secondary antibodies used were a 1:2,000 dilution of donkey anti-rat antibody conjugated with an Alexa-594 fluorophore (Invitrogen, Waltham, MA; Catalog #A-21209) and a 1:2,000 dilution of donkey anti-rabbit antibody conjugated with an Alexa-647 fluorophore (Invitrogen, Waltham, MA; Catalog #A-31573). A confocal microscope (Leica SP8) was used with a 10x objective to acquire images of the entire ipsilateral dorsal horn. Three dorsal horn slices were imaged, then averaged per animal.

#### EphB kinase inhibition

Triple knock-in mice that allow for inducible inhibition of the intracellular kinase activity of EphB1, 2 and 3 in the presence of a PP1 analog were used (referred to as: analog-sensitive EphB triple knock in [AS-EphB TKI] mice).^32^ Mice were administered 1-NA-PP1 dissolved in 10% DMSO/20% Cremaphor-EL/70% saline or the control vehicle alone without 1-NA-PP1. Litter and sex-matched mice were randomly assigned to either the group receiving 1-NA-PP1 or vehicle control. Mice were given intraperitoneal injections every 12 hours (2x per day) for 2 days (behavior analysis) or 5 days (immunohistochemistry).

#### Antibody microarray

The KAM 2000 chip by Kinexus (Vancouver, British Columbia, Canada) was used to evaluate total and phosphorylated protein levels in wild-type and AS-EphB2 TKI mice. For comparing spinal cord injury animals to laminectomy animals, we microdissected the ipsilateral dorsal quadrant of the cervical spinal cord just caudal to the contusion site (i.e., the location of sensory input from primary afferents innervating the forepaw plantar surface) two weeks after surgery. In experiments measuring the effects of EphB1-3 inhibition, we microdissected the ipsilateral C7-C8 dorsal quadrant at the end of the second round of 1-NA-PP1 treatment to generate samples. For each sample used to perform the antibody microarray assay, we pooled dorsal quadrant homogenate from two animals to generate sufficient tissue. Lysate protein from each sample (80-100 μg) was labeled covalently with a fluorescent dye combination. Free dye molecules were then removed via gel filtration. After blocking nonspecific binding sites on the array, an incubation chamber was mounted onto the microarray to permit the loading of the samples. After incubation, unbound proteins were washed away. Two 16-bit images from each array were then captured using a Scan Array Reader (PerkinElmer). The array output consisted of the average normalized net signals (i.e., the average of 4 normalized net signal values of each antibody on the microarray). The standard and percent standard deviations of 4 separate measurements of globally normalized signal intensity values for each antibody on the microarray were calculated. The data are presented as change from control (CFC%). A positive value corresponded to an increase in signal intensity in response to the treatment, with a value of 100% corresponding to a 2-fold increment in signal intensity. A negative CFC value indicated the degree of signal intensity reduction from the controls. Unpaired t-test were used as statistics for the CFC values of the differentially regulated proteins. The CFC values were obtained by gene ontology analysis of the differentially expressed proteins using g-Profiler.

#### Statistical analysis

Behavioral assays were analyzed using a repeated measures two-way ANOVA, and post-hoc comparisons were made both over time and between groups. All qPCR and EphB2 western blot analyses were conducted using a one-way ANOVA across three groups. RNAscope and immunohistochemical analyses were analyzed using a t-test for two groups or ANOVA for three or more groups. Statistics were calculated using GraphPad Prism 9. Antibody microarray analysis was conducted using a generalized linear model under a binomial distribution with logit link function to determine the significance of the abundance among the different categorical groups. All statistical analyses were performed using a package in R (v4.2.1). A p-value was determined with N = 3-4 measurements in each set, paired and 2-tailed in distribution. A p-value of ≤ 0.05 was accepted as statistically significant. Gene ontology analysis was performed using a g:profiler, and the lead proteins with p<0.05 were selected from each group.

## SUPPLEMENTAL FIGURES AND FIGURE LEGENDS

**Supplemental Figure 1.**
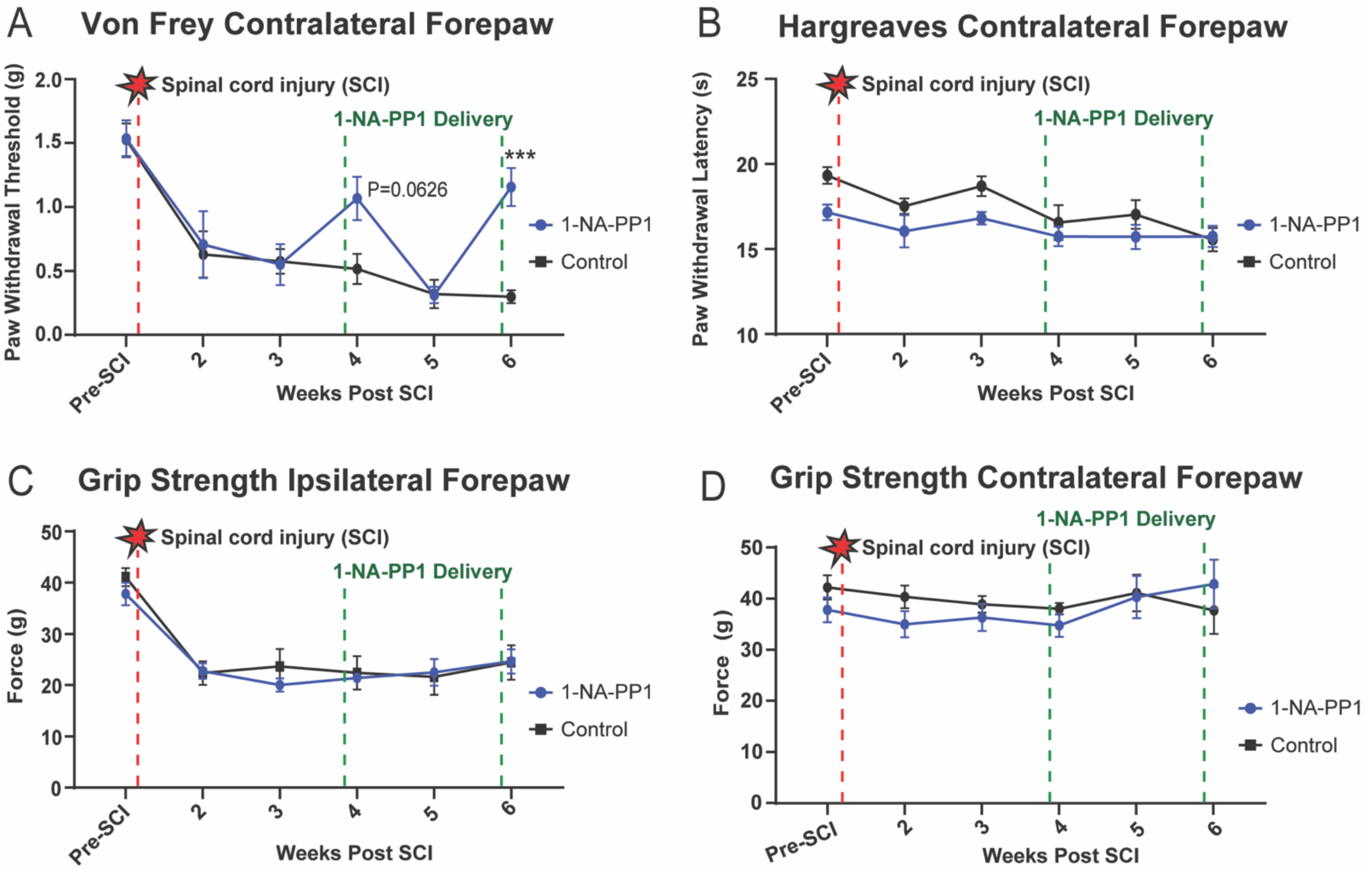
Inducible chemogenetic EphB kinase inhibition reversed mechanical allodynia after spinal cord injury, but did not affect other forelimb sensory or motor behaviors. (A-B) Results of sensory behavioral testing conducted on the left contralesional forepaw, including the (A) Von Frey test and (B) Hargreaves test. All mice received spinal cord injury and were administered either 1-NA-PP1 or vehicle-only control. (C-D) Grip strength testing was performed on spinal cord injury mice administered either 1-NA-PP1 or vehicle-only to assess motor function in each forelimb; no differences in grip strength were measured across groups in either forelimb. Two-way repeated measures ANOVA using Bonferroni’s multiple comparisons were performed between groups to assess significance in all graphs. Significance is shown as * = p<0.05, ** = p<0.01, *** = p<0.001. Detailed statistical analysis for this figure can be found in Supplemental Table 6.

**Supplemental Figure 2.**
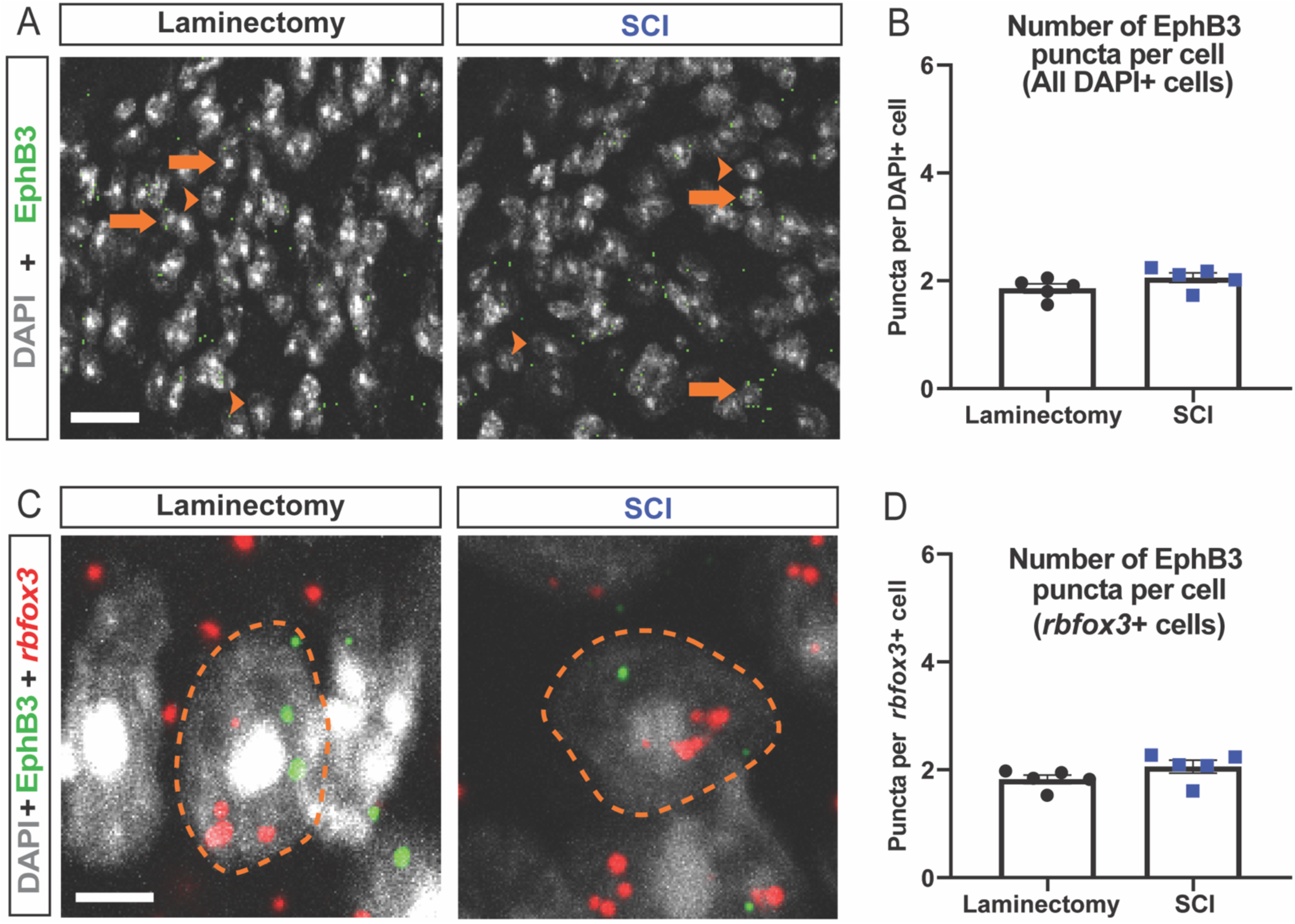
Spinal cord injury did not alter EphB3 expression in cervical dorsal horn neurons. Changes in *Ephb3* expression in superficial laminae of dorsal horn were examined using RNAscope in situ hybridization. (A) *Ephb3* mRNA was counted across all nuclei (marked with DAPI), as shown in representative images from (A, left) laminectomy-only or (A, right) spinal cord injury mice. (A) Arrowheads indicate nuclei without *Ephb3* mRNA, while an arrow indicates *Ephb3-*positive nuclei. (B) Quantification across all nuclei shows no significant increase in *Ephb3* mRNA after spinal cord injury. (C-D) Nuclei were probed for *Rbfox3*, a neuronal marker, along with *Ephb3* mRNA to assess differences in *Ephb3* expression in neurons after either (C, left) laminectomy-only or (C, right) spinal cord injury. (C) Orange dotted lines mark outline of nuclei. Independent (unpaired) samples t-test was used to determine significance between groups in all graphs. * = p<0.05, ** = p<0.01, *** = p<0.001 indicate significance between groups. Detailed statistical analysis for this figure are found in Supplemental Table 7. Scale bars: A (20 mm); C (5 mm).

**Supplemental Figure 3.**
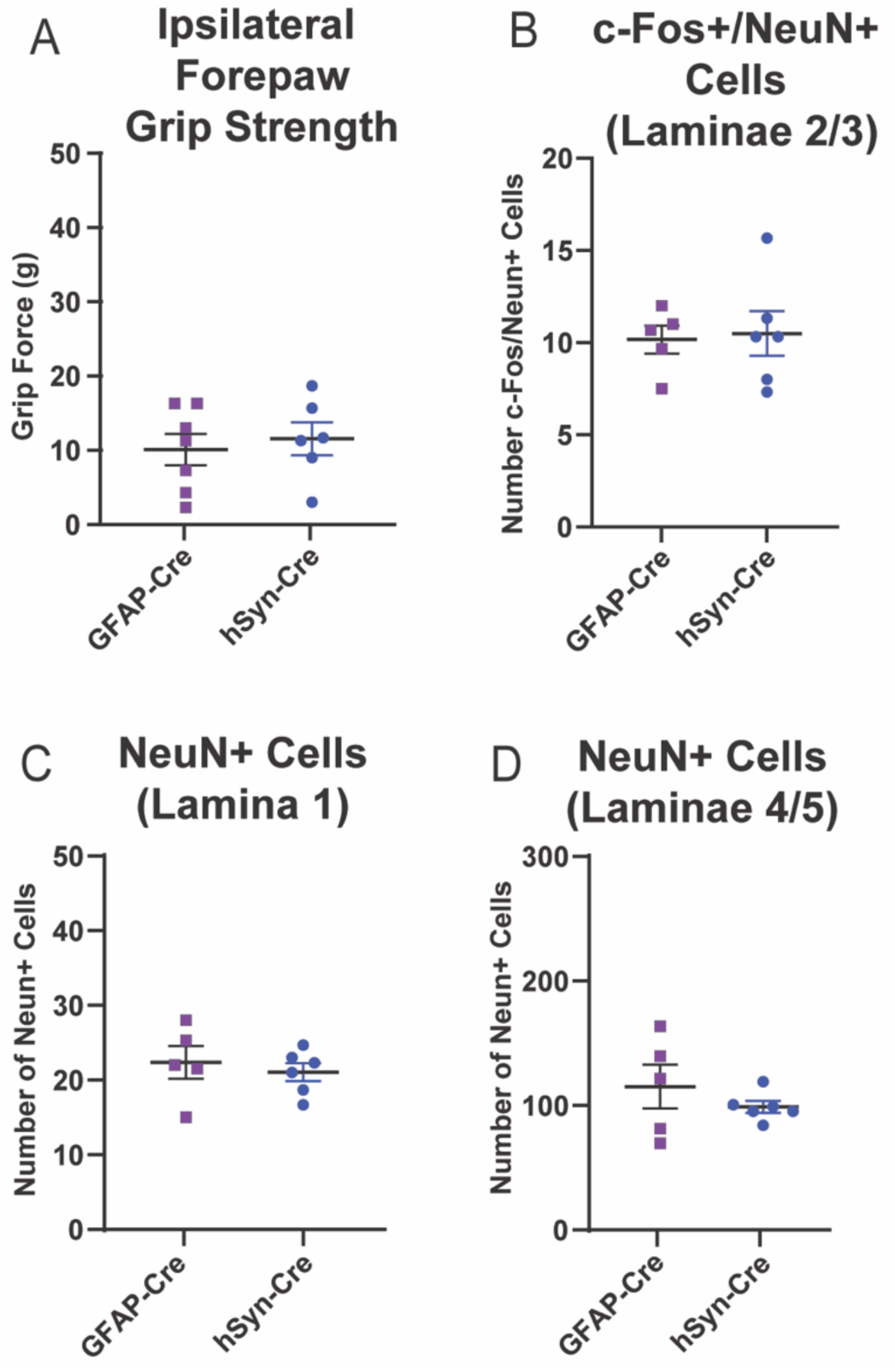
*Ephb2* knockout in neurons or astrocytes of cervical dorsal horn had no effect on forelimb grip strength or dorsal horn neuron numbers. (A) Grip strength was used to assess motor behavior in ipsilesional forepaw. Comparisons were made between mice injected with GFAP-Cre or hSyn-Cre; no significant difference was measured. (B) No difference in number of c-Fos positive neurons (as indicated by NeuN co-labelling) in laminae 2/3 of ipsilateral dorsal horn was observed between GFAP-Cre and hSyn-Cre injected mice. (C, D) No significant difference between virus injection conditions was found in the total number of NeuN-positive neurons in either lamina 1 or laminae 4-5 of ipsilesional dorsal horn. An independent samples t-test was used to determine significance for A and C-F, with * = p<0.05, ** = p<0.01, *** = p<0.001. Detailed statistical analysis for this figure are presented in Supplemental Table 8.

## SUPPLEMENTAL TABLES

**Supplemental Table 1.** Antibodies.

| **Antibody Target** | **Antibody derived from** | **Dilution** | **Source** |
| --- | --- | --- | --- |
| EphB1-pY594 | Rabbit | 1:1,000 | Kinexus, CA |
| EphB3-pY600 | Rabbit | 1:1,000 | Kinexus, CA |
| ERK2-pY263 | Rabbit | 1:1,000 | Kinexus, CA |
| CREBBP-pS121+pS124 | Rabbit | 1:1,000 | Kinexus, CA |
| phospho-Tyrosine (pY99) | Mouse | 1:500 | Santa Cruz Biotechnology, TX |
| EphB2 | Mouse | 1:500 | Invitrogen, MA |
| PKCγ-pT514 | Rabbit | 1:1,000 | Invitrogen, MA |
| Mek1/2 | Rabbit | 1:1,000 | Cell Signaling Technology, MA |
| Tubulin | Rabbit | 1:20,000 | Abcam, UK |

**Supplemental Table 2.** Statistical analysis details for Figure 2.

| **Figure 2 Panel D** |  |  |  |  |  |
| --- | --- | --- | --- | --- | --- |
| Two-way ANOVA<br>1-NA-PP1 vs Control<br>Weeks post Spinal Cord<br>Injury (SCI) | Bonferroni's multiple<br>comparisons test<br>Mean (1-NA-PP1,<br>Control) | N<br>(1-NA-PP1,<br>Control) | Predicted (LS)<br>mean difference | 95.00% CI of<br>difference | Adjusted P<br>Value (< 0.05) |
| Pre-SCI | 1.533, 1.563 | 9, 8 | -0.02917 | -0.4627 to 0.4044 | >0.9999 |
| 2 | 0.4089, 0.3237 | 9, 8 | 0.08514 | -0.3484 to 0.5187 | >0.9999 |
| 3 | 0.3456, 0.2937 | 9, 8 | 0.05181 | -0.3817 to 0.4853 | >0.9999 |
| 4 (1-NA-PP1 Round 1) | 1.067, 0.3137 | 9, 8 | 0.7529 | 0.3194 to 1.186 | **<0.0001** |
| 5 (1 Week wash out) | 0.3056, 0.2837 | 9, 8 | 0.02181 | -0.4117 to 0.4553 | >0.9999 |
| 6 (1-NA-PP1 Round 2) | 1.356, 0.2937 | 9, 8 | 1.062 | 0.6283 to 1.495 | **<0.0001** |
| **Figure 2 Panel E** |  |  |  |  |  |
| Two-way ANOVA<br>Pre-Treatment vs. Post-Treatment | Bonferroni's multiple<br>comparisons test<br>Mean (Pre, Post) | N<br>(Pre-treatment,<br>Post-treatment) | Predicted (LS)<br>mean difference | 95.00% CI of<br>difference | Adjusted P<br>Value (< 0.05) |
| Laminectomy + 1-NA-PP1 | 1.067, 1.267 | 6, 6 | -0.2000 | -0.6961 to 0.2961 | >0.9999 |
| Laminectomy + Control | 1.100, 1.133 | 6, 6 | -0.03333 | -0.5294 to 0.4627 | >0.9999 |
| SCI + 1-NA-PP1 | 0.3456, 1.067 | 9, 9 | -0.7211 | -1.126 to -0.3161 | **0.0003** |
| SCI + Control | 0.2938, 0.3138 | 8, 8 | -0.02000 | -0.4496 to 0.4096 | >0.9999 |
| **Figure 2 Panel F** |  |  |  |  |  |
| Two-way ANOVA<br>1-NA-PP1 vs Control<br>Weeks post SCI | Šídák's multiple<br>comparisons test<br>Mean (1-NA-PP1,<br>Control) | N<br>(1-NA-PP1,<br>Control) | Predicted (LS)<br>mean difference | 95.00% CI of<br>difference | Adjusted P<br>Value (< 0.05) |
| Pre-SCI | 19.51, 18.92 | 9, 8 | 0.5940 | -1.818 to 3.006 | 0.9861 |
| 2 | 15.86, 16.27 | 9, 8 | -0.4111 | -2.824 to 2.001 | 0.9981 |
| 3 | 15.89, 16.88 | 9, 8 | -0.9981 | -3.411 to 1.414 | 0.8470 |
| 4 (1-NA-PP1 Round 1) | 15.61, 15.39 | 9, 8 | 0.2199 | -2.193 to 2.632 | >0.9999 |
| 5 (1 Week wash out) | 15.88, 15.06 | 9, 8 | 0.8190 | -1.593 to 3.231 | 0.9335 |
| 6 (1-NA-PP1 Round 2) | 15.38, 15.13 | 9, 8 | 0.2481 | -2.164 to 2.661 | 0.9999 |
| **Figure 2 Panel G** |  |  |  |  |  |
| Two-way ANOVA<br>Pre-Treatment vs. Post-Treatment | Bonferroni's multiple<br>comparisons test<br>Mean (Pre, Post) | N<br>(Pre-treatment,<br>Post-treatment) | Predicted (LS)<br>mean difference | 95.00% CI of<br>difference | Adjusted P<br>Value (< 0.05) |
| Laminectomy + 1-NA-PP1 | 19.01, 17.97 | 6, 6 | 1.039 | -1.850 to 3.928 | 0.8146 |
| Laminectomy + Control | 19.45, 18.96 | 6, 6 | 0.4972 | -2.391 to 3.386 | 0.9847 |
| SCI + 1-NA-PP1 | 15.89, 15.62 | 9, 9 | 0.2630 | -2.096 to 2.622 | 0.9971 |
| SCI + Control | 16.88, 15.39 | 8, 8 | 1.496 | -1.006 to 3.997 | 0.4035 |

**Supplemental Table 3.** Statistical analysis details for Figure 3.

| Figure 3 Panel B |  |  |  |  |  |
| --- | --- | --- | --- | --- | --- |
| One-way ANOVA<br>Tukey's multiple comparisons test | Mean | N | Mean difference | 95.00% CI of difference | Adjusted P Value (< 0.05) |
| Laminectomy vs. 40 kD SCI | 1.016, 0.9951 | 8, 8 | 0.02051 | -0.2882 to 0.3292 | 0.9847 |
| Laminectomy vs. 60 kD SCI | 1.016, 0.9428 | 8, 8 | 0.07280 | -0.2359 to 0.3815 | 0.8246 |
| 40 kD SCI vs. 60 kD SCI | 0.9951, 0.9428 | 8, 8 | 0.05230 | -0.2564 to 0.3610 | 0.9048 |
| Figure 3 Panel C |  |  |  |  |  |
| One-way ANOVA<br>Tukey's multiple comparisons test |  | N | Mean difference | 95.00% CI of difference | Adjusted P Value (< 0.05) |
| Laminectomy vs. 40 kD SCI | 1.016, 1.285 | 8, 8 | -0.2694 | -0.5355 to -0.003296 | 0.0469 |
| Laminectomy vs. 60 kD SCI | 1.016, 1.545 | 8, 8 | -0.5285 | -0.7946 to -0.2624 | 0.0002 |
| 40 kD SCI vs. 60 kD SCI | 1.285, 1.545 | 8, 8 | -0.2591 | -0.5252 to 0.006979 | 0.0572 |
| Figure 3 Panel D |  |  |  |  |  |
| One-way ANOVA<br>Tukey's multiple comparisons test |  | N | Mean difference | 95.00% CI of difference | Adjusted P Value (< 0.05) |
| Laminectomy vs. 40 kD SCI | 1.021, 1.243 | 8, 8 | -0.2220 | -0.4658 to 0.02176 | 0.0785 |
| Laminectomy vs. 60 kD SCI | 1.021, 1.380 | 8, 8 | -0.3587 | -0.6025 to -0.1149 | 0.0036 |
| 40 kD SCI vs. 60 kD SCI | 1.243, 1.380 | 8, 8 | -0.1367 | -0.3805 to 0.1071 | 0.3525 |
| Figure 3 Panel H |  |  |  |  |  |
| Unpaired t-test<br>Laminectomy vs. SCI<br>Mean | Difference Between Means ± (SEM) |  | N | 95% Confidence Interval | P value (< 0.05) |
| 4.601, 5.101 | 0.4994 ± 0.2265 |  | 7, 7 | 0.0059 to 0.9928 | 0.0477 |
| Figure 3 Panel J |  |  |  |  |  |
| Unpaired t-test<br>Laminectomy vs. SCI<br>Mean | Difference Between Means ± (SEM) |  | N | 95% Confidence Interval | P value (< 0.05) |
| 6.894, 7.942 | 1.048 ± 0.2200 |  | 7, 7 | 0.5689 to 1.528 | 0.0005 |
| Figure 3 Panel L |  |  |  |  |  |
| Unpaired t-test<br>Laminectomy vs. SCI<br>Mean | Difference Between Means ± (SEM) |  | N | 95% Confidence Interval | P value (< 0.05) |
| 3.250, 4.067 | 0.8168 ± 0.3614 |  | 7, 7 | 0.02947 to 1.604 | 0.0432 |

**Supplemental Table 4.** Statistical analysis details for Figure 4.

| Figure 4 Panel D |  |  |  |  |  |
| --- | --- | --- | --- | --- | --- |
| Two-way ANOVA<br>hSyn-Cre vs. GFAP-Cre<br>Weeks post SCI | Bonferroni's multiple<br>comparisons test<br>Mean (hSyn, GFAP) | N<br>(hSyn-Cre,<br>GFAP-Cre) | Predicted (LS)<br>mean difference | 95.00% CI of<br>difference | Adjusted P<br>Value (< 0.05) |
| Pre-SCI | 1.871, 1.850 | 7, 6 | 0.02143 | -0.5598 to 0.6026 | >0.9999 |
| 3 | 0.8286, 0.6667 | 7, 6 | 0.1619 | -0.4193 to 0.7431 | 0.9731 |
| 4 | 0.7714, 0.7333 | 7, 6 | 0.03810 | -0.5431 to 0.6193 | >0.9999 |
| 7 (2 Weeks Post AAV) | 1.771, 0.7000 | 7, 6 | 1.071 | 0.4902 to 1.653 | <0.0001 |
| 9 (4 Weeks Post AAV) | 1.657, 0.9667 | 7, 6 | 0.6905 | 0.1093 to 1.272 | 0.0118 |
| 11 (6 Weeks Post AAV) | 1.486, 0.6333 | 7, 6 | 0.8524 | 0.2712 to 1.434 | 0.0011 |
| Two-way ANOVA<br>hSyn-Cre vs. hSyn-GFP<br>Weeks post SCI | Bonferroni's multiple<br>comparisons test<br>Mean (hSyn, GFAP) | N<br>(hSyn-Cre,<br>GFAP-Cre) | Predicted (LS)<br>mean difference | 95.00% CI of<br>difference | Adjusted P<br>Value (< 0.05) |
| Pre-SCI | 1.871, 1.822 | 7, 9 | 0.04921 | -0.4586 to 0.5571 | >0.9999 |
| 3 | 0.8286, 0.6444 | 7, 9 | 0.1841 | -0.3237 to 0.6920 | 0.9966 |
| 4 | 0.7714, 0.7778 | 7, 9 | -0.006349 | -0.5142 to 0.5015 | >0.9999 |
| 7 (2 Weeks Post AAV) | 1.771, 0.5067 | 7, 9 | 1.265 | 0.7569 to 1.773 | <0.0001 |
| 9 (4 Weeks Post AAV) | 1.657, 0.6889 | 7, 9 | 0.9683 | 0.4604 to 1.476 | <0.0001 |
| 11 (6 Weeks Post AAV) | 1.486, 0.7778 | 7, 9 | 0.2778 | 0.2001 to 1.216 | 0.0008 |
| Two-way ANOVA<br>GFAP-Cre vs. hSyn-GFP<br>Weeks post SCI | Bonferroni's multiple<br>comparisons test<br>Mean (hSyn, GFAP) | N<br>(hSyn-Cre,<br>GFAP-Cre) | Predicted (LS)<br>mean difference | 95.00% CI of<br>difference | Adjusted P<br>Value (< 0.05) |
| Pre-SCI | 1.850, 1.822 | 6, 9 | 0.02778 | -0.5033 to 0.5589 | >0.9999 |
| 3 | 0.6667, 0.6444 | 6, 9 | 0.02222 | -0.5089 to 0.5533 | >0.9999 |
| 4 | 0.7333, 0.7778 | 6, 9 | -0.04444 | -0.5756 to 0.4867 | >0.9999 |
| 7 (2 Weeks Post AAV) | 0.7000, 0.5067 | 6, 9 | 0.1933 | -0.3378 to 0.7244 | 0.9965 |
| 9 (4 Weeks Post AAV) | 0.9667, 0.6889 | 6, 9 | 0.7079 | -0.2533 to 0.8089 | 0.8855 |
| 11 (6 Weeks Post AAV) | 0.6333, 0.7778 | 6, 9 | -0.1444 | -0.6756 to 0.3867 | >0.9999 |
| Figure 4 Panel F |  |  |  |  |  |
| Unpaired t-test<br>GFAP-Cre vs. hSyn-Cre<br>Mean (GFAP, hSyn) | Difference Between Means ± (SEM) |  | N | 95% Confidence Interval | P value<br>(< 0.05) |
| 5.833, 3.889 | -1.944 ± 0.5762 |  | 5, 6 | -3.248 to -0.6411 | 0.0082 |
| Figure 4 Panel G |  |  |  |  |  |
| Unpaired t-test<br>GFAP-Cre vs. hSyn-Cre<br>Mean (GFAP, hSyn) | Difference Between Means ± (SEM) |  | N | 95% Confidence Interval | P value<br>(< 0.05) |
| 22.57, 14.67 | -7.900 ± 3.146 |  | 5, 6 | -15.02 to -0.7842 | 0.0332 |

**Supplemental Table 5.** Statistical analysis details for Supplemental Figure 1.

| **Supplemental Figure 1 Panel A** |  |  |  |  |  |
| --- | --- | --- | --- | --- | --- |
| Two-way ANOVA<br>1-NA-PP1 vs Control<br>Weeks post SCI | Šídák's multiple<br>comparisons test<br>Mean (1-NA-PP1,<br>Control) | N<br>(1-NA-PP1,<br>Control) | Predicted (LS)<br>mean difference | 95.00% CI of<br>difference | Adjusted P<br>Value (< 0.05) |
| Pre-SCI | 1.533, 1.525 | 9, 8 | 0.008333 | -0.5612 to 0.5778 | >0.9999 |
| 2 | 0.7056, 0.6287 | 9, 8 | 0.07681 | -0.4927 to 0.6463 | 0.9995 |
| 3 | 0.5500, 0.5750 | 9, 8 | -0.02500 | -0.5945 to 0.5445 | >0.9999 |
| 4 (1-NA-PP1 Round 1) | 1.067, 0.5150 | 9, 8 | 0.5517 | -0.01784 to 1.121 | 0.0626 |
| 5 (1 Week wash out) | 0.3111, 0.3200 | 9, 8 | -0.008889 | -0.5784 to 0.5606 | >0.9999 |
| 6 (1-NA-PP1 Round 2) | 1.156, 0.2987 | 9, 8 | 0.8568 | 0.2873 to 1.426 | **0.0007** |
| **Supplemental Figure 1 Panel B** |  |  |  |  |  |
| Two-way ANOVA<br>1-NA-PP1 vs Control<br>Weeks post SCI | Šídák's multiple<br>comparisons test<br>Mean (1-NA-PP1,<br>Control) | N<br>(1-NA-PP1,<br>Control) | Predicted (LS)<br>mean difference | 95.00% CI of<br>difference | Adjusted P<br>Value (< 0.05) |
| Pre-SCI | 17.15, 19.32 | 9, 8 | -2.167 | -4.725 to 0.3911 | 0.1412 |
| 2 | 16.04, 17.52 | 9, 8 | -1.472 | -4.030 to 1.086 | 0.5514 |
| 3 | 16.81, 18.69 | 9, 8 | -1.880 | -4.438 to 0.6779 | 0.2699 |
| 4 (1-NA-PP1 Round 1) | 15.74, 16.55 | 9, 8 | -0.8171 | -3.375 to 1.741 | 0.9497 |
| 5 (1 Week wash out) | 15.71, 17.03 | 9, 8 | -1.314 | -3.872 to 1.244 | 0.6742 |
| 6 (1-NA-PP1 Round 2) | 15.73, 15.54 | 9, 8 | 0.1917 | -2.366 to 2.750 | >0.9999 |
| **Supplemental Figure 1 Panel C** |  |  |  |  |  |
| Two-way ANOVA<br>1-NA-PP1 vs Control<br>Weeks post SCI | Šídák's multiple<br>comparisons test<br>Mean (1-NA-PP1,<br>Control) | N<br>(1-NA-PP1,<br>Control) | Predicted (LS)<br>mean difference | 95.00% CI of<br>difference | Adjusted P<br>Value (< 0.05) |
| Pre-SCI | 37.78, 41.08 | 9, 8 | -3.306 | -12.66 to 6.044 | 0.9204 |
| 2 | 22.74, 22.33 | 9, 8 | 0.4074 | -8.942 to 9.757 | >0.9999 |
| 3 | 20.00, 23.62 | 9, 8 | -3.625 | -12.97 to 5.725 | 0.8821 |
| 4 (1-NA-PP1 Round 1) | 21.41, 22.37 | 9, 8 | -0.9676 | -10.32 to 8.382 | 0.9999 |
| 5 (1 Week wash out) | 22.48, 21.58 | 9, 8 | 0.8981 | -8.452 to 10.25 | >0.9999 |
| 6 (1-NA-PP1 Round 2) | 24.63, 24.42 | 9, 8 | 0.2130 | -9.137 to 9.563 | >0.9999 |
| **Supplemental Figure 1 Panel D** |  |  |  |  |  |
| Two-way ANOVA<br>1-NA-PP1 vs Control<br>Weeks post SCI | Šídák's multiple<br>comparisons test<br>Mean (1-NA-PP1,<br>Control) | N<br>(1-NA-PP1,<br>Control) | Predicted (LS)<br>mean difference | 95.00% CI of<br>difference | Adjusted P<br>Value (< 0.05) |
| Pre-SCI | 37.80, 42.19 | 9, 8 | -4.391 | -16.19 to 7.404 | 0.9005 |
| 2 | 34.96, 40.33 | 9, 8 | -5.370 | -17.17 to 6.425 | 0.7814 |
| 3 | 36.30, 38.88 | 9, 8 | -2.579 | -14.37 to 9.217 | 0.9925 |
| 4 (1-NA-PP1 Round 1) | 34.74, 38.00 | 9, 8 | -3.259 | -15.05 to 8.536 | 0.9750 |
| 5 (1 Week wash out) | 40.30, 41.08 | 9, 8 | -0.7870 | -12.58 to 11.01 | >0.9999 |
| 6 (1-NA-PP1 Round 2) | 42.85, 37.67 | 9, 8 | 5.185 | -6.610 to 16.98 | 0.8074 |

**Supplemental Table 6.** Statistical analysis details for Supplemental Figure 2.

| **Supplemental Figure 2 Panel B** |  |  |  |  |
| --- | --- | --- | --- | --- |
| Unpaired t-test<br>Laminectomy vs. SCI<br>Mean (Lam., SCI) | Difference Between<br>Means ± (SEM) | N | 95% Confidence Interval | P value<br>(**< 0.05**) |
| 1.858, 2.057 | 0.1997 ± 0.1231 | 5, 5 | -0.08422 to 0.4837 | 0.1435 |
| **Supplemental Figure 2 Panel D** |  |  |  |  |
| Unpaired t-test<br>Laminectomy vs. SCI<br>Mean (Lam., SCI) | Difference Between<br>Means ± (SEM) | N | 95% Confidence Interval | P value<br>(**< 0.05**) |
| 1.823, 2.058 | 0.2347 ± 0.1432 | 5, 5 | -0.09539 to 0.5648 | 0.1397 |

**Supplemental Table 7.** Statistical analysis details for Supplemental Figure 3.

| **Supplemental Figure 3 Panel A** |  |  |  |  |
| --- | --- | --- | --- | --- |
| Unpaired t-test<br>GFAP-Cre vs. hSyn-Cre<br>Mean (GFAP, hSyn) | Difference Between Means<br>± (SEM) | N | 95% Confidence Interval | P value<br>(**< 0.05**) |
| 10.11, 11.55 | 1.437 ± 3.077 | 6, 7 | -8.209 to 5.334 | 0.6495 |
| **Supplemental Figure 3 Panel B** |  |  |  |  |
| Unpaired t-test<br>GFAP-Cre vs. hSyn-Cre<br>Mean (GFAP, hSyn) | Difference Between Means<br>± (SEM) | N | 95% Confidence Interval | P value<br>(**< 0.05**) |
| 10.17, 10.50 | 0.3333 ± 1.502 | 5, 6 | -3.065 to 3.732 | 0.8293 |
| **Supplemental Figure 3 Panel C** |  |  |  |  |
| Unpaired t-test<br>GFAP-Cre vs. hSyn-Cre<br>Mean (GFAP, hSyn) | Difference Between Means<br>± (SEM) | N | 95% Confidence Interval | P value<br>(**< 0.05**) |
| 22.37, 21.06 | -1.311 ± 2.381 | 5, 6 | -6.697 to 4.074 | 0.5952 |
| **Supplemental Figure 3 Panel D** |  |  |  |  |
| Unpaired t-test<br>GFAP-Cre vs. hSyn-Cre<br>Mean (GFAP, hSyn) | Difference Between Means<br>± (SEM) | N | 95% Confidence Interval | P value<br>(**< 0.05**) |
| 115.2, 98.89 | -16.28 ± 16.71 | 5, 6 | -54.09 to 21.53 | 0.3555 |

